# Granulosa cell genes that regulate ovarian follicle development beyond the antral stage: the role of estrogen receptor β

**DOI:** 10.1101/2020.09.25.295550

**Authors:** V. Praveen Chakravarthi, Shaon Borosha, Subhra Ghosh, Katherine F. Roby, Michael W. Wolfe, Lane K. Christenson, M. A. Karim Rumi

## Abstract

Follicle development beyond the preantral stage is dependent on gonadotropins. FSH signaling is crucial for the advancement of preantral follicles to the antral stage, and LH signaling is essential for further maturation of preovulatory follicles. Estrogen is intricately tied to gonadotropin signaling during the advanced stages of folliculogenesis. We observed that *Erβ*^*null*^ ovarian follicles fail to develop beyond the antral stage, even after exogenous gonadotropin stimulation. As ERβ is primarily expressed in the granulosa cells (GCs), we explored the gonadotropin-regulated GC genes that induce maturation of antral follicles. Synchronized follicle development was induced by administration of exogenous gonadotropins to wildtype 4-wk-old female rats. The GC transcriptome was analyzed via RNA-sequencing before and after gonadotropin stimulation. An *Erβ*^*null*^ mutant model that fails to show follicle maturation was also included in order to identify the ERβ-regulated genes involved at this step. We observed that specific groups of genes were differentially expressed in response to PMSG or hCG administration in wildtype rats. While some of the PMSG or hCG-induced genes showed a similar expression pattern in *Erβ*^*null*^ GCs, a subset of PMSG- or hCG-induced genes showed a differential expression in *Erβ*^*null*^ GCs. These latter ERβ-regulated genes included previously known FSH or LH target genes including *Lhcgr, Cyp11a1, Cyp19a1, Pgr, Runx2, Egfr, Kiss1*, and *Ptgs2*, which are involved in follicle development, oocyte maturation, and ovulation. We also identified novel ERβ-regulated genes including *Jaml, Galnt6, Znf750, Dusp9, Wnt16*, and *Mageb16* that failed to respond to gonadotropin stimulation in *Erβ*^*null*^ GCs. Our findings indicate that the gonadotropin-induced spatiotemporal pattern of gene expression is essential for ovarian follicle maturation beyond the antral stage. However, expression of a subset of those gonadotropin-induced genes is dependent on transcriptional regulation by ERβ.

## 1. INTRODUCTION

Ovarian granulosa (GCs) and theca cells (TCs) play essential regulatory roles in follicle development and oocyte maturation (Eppig, 2001; Jones and Shikanov, 2019; Orisaka et al., 2009). While ovarian follicle activation is gonadotropin-independent, later stages of follicle development require follicle stimulating hormone (FSH) and luteinizing hormone (LH) to induce maturation and ovulation (Goldman and Mahesh, 1968; Goldman and Mahesh, 1969; Grimek et al., 1976; Schwartz, 1974). Gonadotropin signaling initiates steroidogenesis in TCs, which produce androgens (Fortune and Armstrong, 1977) that are converted into estrogens by the neighboring GCs (Erickson and Hsueh, 1978; Liu and Hsueh, 1986). Follicular estrogen in turn regulates the gonadotropin secretion from the H-P axis (Yasin et al., 1995). Gonadotropins also induce differentiation of GCs, which is essential for the expression of factors that regulate the final stages of follicle development, oocyte maturation, and ovulation (Greep et al., 1942; LOSTROH and JOHNSON, 1966). FSH acts through binding the FSH receptor (FSHR) expressed exclusively on GCs, whereas LH acts through binding the LH receptor (LHCGR) expressed on both TCs and GCs (Gromoll et al., 1996; Moudgal, 2012; RAJANIEMI et al., 1977; Simoni et al., 1997). The expression of FSHR on GCs is follicular-stage-specific (Oktay et al., 1997; Peters, 1979), however, the expression of LHCGR on GCs is induced by FSH signaling (Dierich et al., 1998; Zeleznik et al., 1974).

Estrogen signaling plays a dynamic role in ovarian folliculogenesis and steroidogenesis, as well as in the hypothalamic-pituitary-ovarian (H-P-O) axis’ regulation of ovarian functions (Richards, 1980; Shupnik, 2002; Wójtowicz et al., 2007). While TCs predominantly express estrogen receptor alpha (ERα), GCs primarily express estrogen receptor beta (ERβ). Mutant mouse and rat models homozygous for the *Erβ* null mutation suffer from defective follicle maturation and failure of ovulation (Couse et al., 2005a; Rumi et al., 2017). Although *Erβ* mutants exhibit an attenuated gonadotropin surge (Jayes et al., 2014; Rumi et al., 2017), administration of exogenous gonadotropins fails to compensate for the defective follicle maturation and induction of ovulation, which suggests a primary intraovarian defect(s) caused by the loss of ERβ (Rumi et al., 2017). Due to the predominant expression of ERβ in GCs, we suspect that dysregulation of ERβ-regulated genes in GCs is responsible for the failure of gonadotropin-induced follicle maturation.

Gonadotropin stimulation induces the differentiation of somatic cells in the ovary and regulates the expression of genes that impact follicle development and oocyte maturation (Dekel and Beers, 1978; Gómez et al., 1993). We observed that *Erβ*^*null*^ ovarian follicles fail to develop beyond the antral stage. Previous studies have focused on the attenuated gonadotropin surge and defective expression of GC genes involved in the preovulatory process (Khristi et al., 2018; Rumi et al., 2017). However, ERβ is the predominant estrogen receptor expressed in the ovary since the embryonic stage (Byers et al., 1997b; Couse et al., 2004a; Pelletier and El-Alfy, 2000a; Slomczynska et al., 2001). Thus, in the absence of ERβ, ovarian follicle defects may develop due to alerted gene expression prior to gonadotropin stimulation. Therefore, in this study, we have analyzed the GC transcriptome in *Erβ*^*null*^ rat ovaries before and after gonadotropin stimulation and compared them with those of wildtype rats. This systematic approach generated an invaluable source of data understanding the molecular mechanisms involved in ERβ-regulated follicle maturation beyond the antral stage.

## 2. MATERIALS AND METHODS

### 2.1. Animal models

4-wk-old *Erβ*^*null*^ and age-matched wildtype female Holtzman Sprague-Dawley (HSD) rats were used in this study. *Erβ*^*null*^ mutant rats were generated by targeted deletion of exon 3 of the rat *Erβ* gene (Rumi et al., 2017). Experimental animals were generated by breeding heterozygous mutant male and female rats carrying the exon 3 deletion within *Erβ* gene. Presence of the *Erβ* gene mutation was screened for using PCR-based genotyping (Rumi et al., 2017). All animal experiments were performed in accordance with the protocols approved by the University of Kansas Medical Center (KUMC) Animal Care and Use Committee.

### 2.2. Induction of follicle maturation by gonadotropin administration

Synchronized follicular growth was initiated by administration of exogenous gonadotropins to 4-wk-old *Erβ*^*null*^ and wildtype female rats as described previously by our laboratory (Chakravarthi et al., 2020a; Chakravarthi et al., 2018; Khristi et al., 2018; Rumi et al., 2017). Briefly, 30IU of pregnant mare’s serum gonadotropin (PMSG, BioVendor, MO) was injected i.p. into immature female rats, and 48h later some of PMSG treated females received an i.p. injection of 30IU of human chorionic gonadotropin (hCG, BioVendor, MO; **Fig. 2A**).

### 2.3. Collection of GCs and RNA-sequencing

Ovarian tissues were collected prior to gonadotropin administration (Basal group), 48h after PMSG injection (PMSG group), and in PMSG-treated rats 4h following hCG injection and 10h post-hCG (hCG 4h and hCG 10h groups). GCs were released from the ovaries by needle puncture under microscopic examination, followed by filtration with a 40μM cell strainer (Chakravarthi et al., 2018; Khristi et al., 2018). Total RNA was extracted from the GCs using TRI Reagent (Sigma-Aldrich, St. Louis, MO) following the manufacturer’s instructions. RNA quality was assessed on an Agilent Bioanalyzer, and samples with RIN values over 9 were selected for mRNA-seq library preparation. Each library was prepared using RNA samples from 3 individual rats and 3 independent libraries were made for each experimental time point. 500 ng of total RNA was used for each RNA-seq library preparation using a TruSeq Standard mRNA kit (Illumina, San Diego, CA) as described in our previous publications (Khristi et al., 2018; Khristi et al., 2019). The cDNA libraries were first evaluated for quality at the KUMC Genomics Core and then sequenced on an Illumina HiSeq X sequencer (Novogene Corporation, Sacramento, CA). All RNA-seq data have been submitted to the Sequencing Read Archive as shown in **Supplementary Table 1**. Additionally, our previously published RNA-seq data of wildtype and *Erβ*^*null*^ GCs collected 10h post-hCG can be used for comparisons (Khristi et al., 2018).

### 2.4. Analyses of RNA-sequencing data

RNA-sequencing data were demultiplexed, trimmed, aligned, and analyzed using CLC Genomics Workbench 20 (Qiagen Bioinformatics, Germantown, MD) as described previously (Chakravarthi et al., 2020a; Chakravarthi et al., 2019; Khristi et al., 2019). Trimming was performed to remove low quality reads, and good quality reads were aligned with the *Rattus norvegicus* genome (Rnor_6.0) using the default parameters: (a) maximum number of allowable mismatches was 2; (b) minimum length and similarity fraction was set at 0.8; and (c) minimum number of hits per read was 10. Expression values were measured in TPM (transcripts per million). The threshold *p*-value was determined according to the false discovery rate (FDR). Differentially expressed genes (DEGs) were selected if the absolute fold change in expression was ≥ 2 with an FDR *p*-value of ≤0.05 as described previously (Chakravarthi et al., 2020a; Chakravarthi et al., 2019; Khristi et al., 2019).

### 2.5. Pathway analyses

In the first approach, DEGs were identified between the ‘Basal to PMSG’ and ‘PMSG to hCG 4h’ groups for either wildtype or *Erβ*^*null*^ GCs. In the second approach, DEGs were identified in Basal, PMSG, or hCG 4h groups comparing the wildtype and *Erβ*^*null*^ data. These DEGs were sub-grouped into upregulated and downregulated genes. Using the Meta-Chart (https://www.meta-chart.com/) software, Venn diagrams were generated by overlapping the upregulated or downregulated DEGs in wildtype and *Erβ*^*null*^ data as identified in the first approach. For further analyses, these Venn diagrams were overlapped with genes upregulated or downregulated in *Erβ*^*null*^ GCs as identified in the second approach. PMSG- or hCG-regulated genes in wildtype GCs that showed a differential expression in *Erβ*^*null*^ GCs were selected as ERβ-regulated genes and subjected to Ingenuity Pathway Analysis (IPA; Qiagen Bioinformatics) to determine their involvement in folliculogenesis.

### 2.6. Validation of RNA-sequencing by RT-qPCR

For validation of RNA-seq data, selected DEGs were analyzed using RT-qPCR. cDNA was prepared from 1 μg of total RNA purified from the GCs collected at the indicated time points (Basal, PMSG, hCG 4h and hCG 10h) (**Fig. 2A**) using the High Capacity cDNA Reverse Transcription kit (ThermoFisher Scientific). qPCR was performed using Power SYBR Green Master Mix (ThermoFisher Scientific). A list of qPCR primer sequences used in this study is shown in **Supplementary Table 2**. The results of RT-qPCR were normalized to *Rn18s* expression and calculated by the comparative ΔΔCT method (Chakravarthi et al., 2020b; Khristi et al., 2018; Khristi et al., 2019).

### 2.7. Statistical analyses

Each experimental group consisted of a minimum of 6 rats, and all procedures were repeated for reproducibility (except for RNA-sequencing). The experimental results are presented as the mean ± standard error (SE). The results were analyzed for one-way ANOVA, and the significance of mean differences was determined by Duncan’s post hoc test, with *p*< 0.05. All the statistical calculations were done using SPSS 22 (IBM, Armonk, NY). Each RNA-seq library was prepared using pooled RNA samples from 3 individual wildtype or *Erβ*^*null*^ rats. Each group of RNA-sequencing data consisted of three different libraries.

## 3. RESULTS

### 3.1. Defective follicle development in *Erβ*^*null*^ rats

Wildtype rat ovaries responded to exogenous gonadotropins and follicles were observed to have varying stages of development including preantral, antral, and large Graafian follicles **(Fig. 1A-C**). In contrast, gonadotropin-stimulated *Erβ*^null^ rat ovaries exhibited an abundance of antral follicles of similar size (**Fig. 1D-F**). While the follicles in wildtype ovaries showed the signs of selection and dominance in response to gonadotropin stimulation (**Fig. 1G**), the antral follicles in *Erβ*^null^ ovaries failed to undergo such preovulatory development to Graafian follicles (**Fig. 1H**).

**Figure 1.**
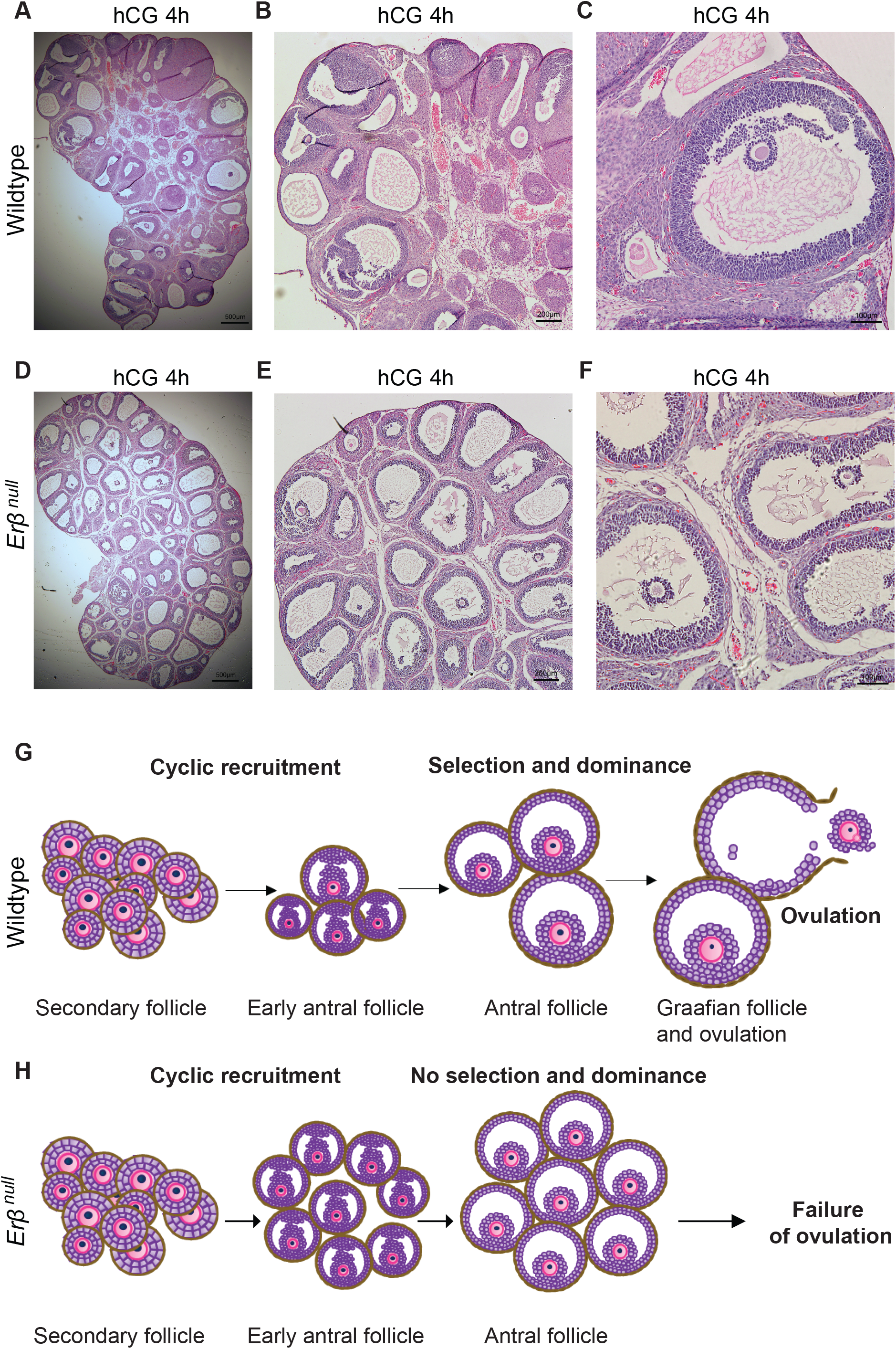
Defective follicle maturation in *Erβ*^*null*^ ovaries. Hematoxylin and Eosin stained sections of ovaries collected from gonadotropin-treated 4-wk-old wildtype rats show different stages of follicles including preantral, early antral, antral, and preovulatory Graafian follicles (**A-C**). In contrast, the sections of ovaries collected from *Erβ*^*null*^ rats show numerous antral follicles of similar size (**D-F**). *Erβ*^*null*^ follicles do not undergo selection and dominance in response to gonadotropin and lack preovulatory maturation, which is observed in wildtype ovaries (**G** and **H**). hCG 4h, 4h after hCG administration to PMSG treated rats.

**Figure 2.**
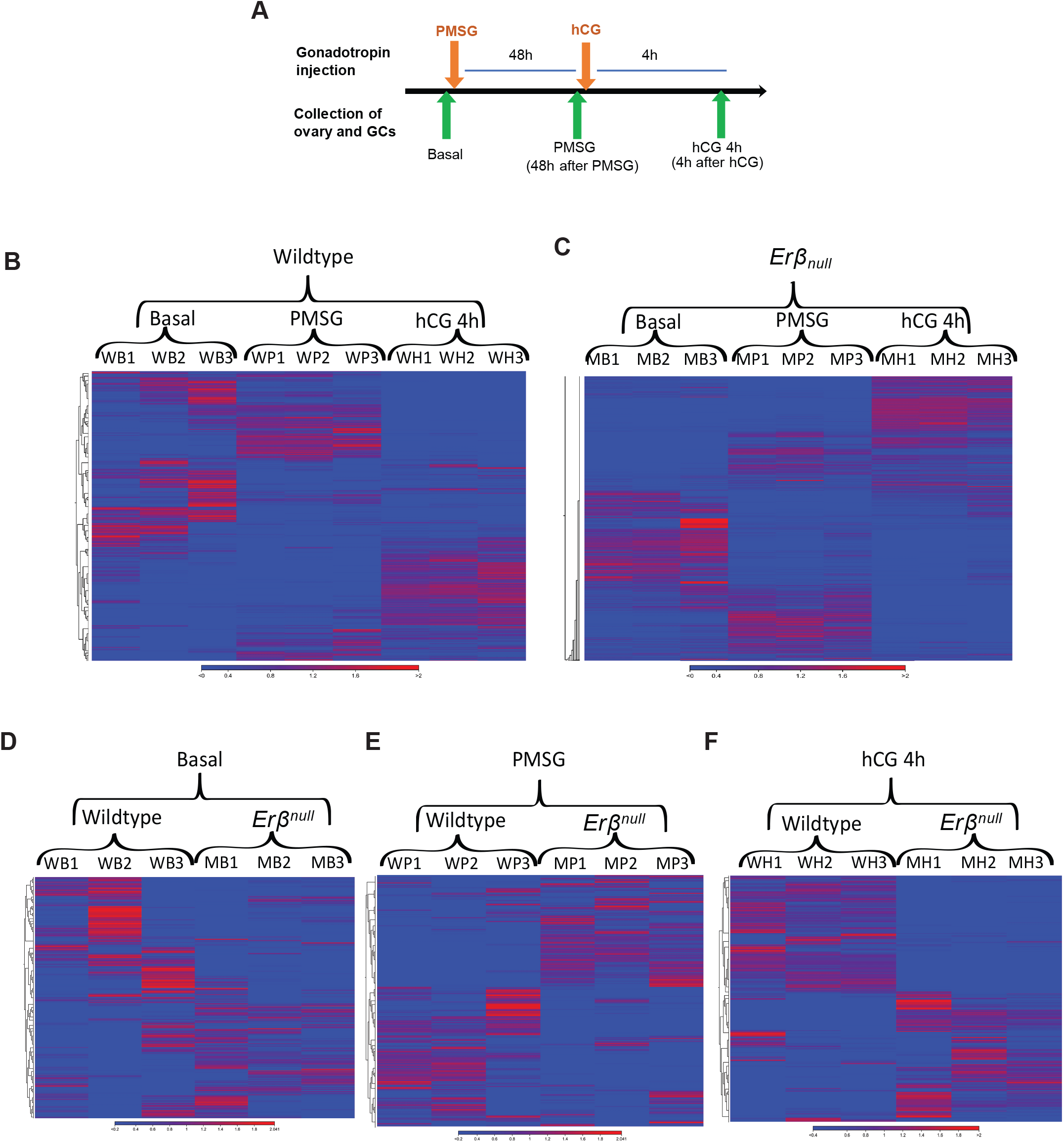
RNA-sequencing of granulosa cells collected from gonadotropin-treated wildtype and *Erβ*^*null*^ rat ovaries. 4-wk-old wildtype and *Erβ*^*null*^ rats were treated with gonadotropins, and ovaries were collected before and after gonadotropin treatment for isolation of granulosa cells (GCs): before treatment (Basal), 48 hours after PMSG injection (PMSG), and 4h after hCG injection (hCG 4h) (**A**). Sequencing was performed on total RNA extracted from GCs, and data were analyzed by using CLC Genomics Workbench. Hierarchical clustering was performed on the differentially expressed genes (DEGs) among replicates at different stages of gonadotropin treatment in wildtype (**B**) or *Erβ*^*null*^ (**C**) GCs. In addition, clustering was performed on the DEGs between wildtype and *Erβ*^*null*^ rat GCs in Basal (**D**), PMSG (**E**), and hCG 4h (**F**) groups.

### 3.2. GC-transcriptome during gonadotropin-induced follicle development

GC mRNA-seq data for Basal, PMSG, and 4h post-hCG groups were analyzed in two different approaches. In the first approach, DEGs were identified across the treatment groups from Basal to PMSG, and from PMSG to 4h post-hCG within the same genotype (i.e., wildtype or *Erβ*^null^; **Fig. 2B** and **2C**). PMSG administration resulted in differential expression of 1720 genes in wildtype and 1085 genes in *Erβ*^null^ GCs. At 4h post-hCG, 3061 genes were differentially expressed in wildtype and 1055 genes in *Erβ*^null^ GCs. The PMSG and 4h post-hCG DEGs in wildtype and *Erβ*^null^ GCs are presented in **Supplementary Tables 3-6**. In the second approach, DEGs were identified within the Basal, PMSG, or 4h post-hCG groups across the wildtype and *Erβ*^null^ RNA-seq data (**Fig. 2D-F**). In in *Erβ*^null^ GCs, 970 genes were differentially expressed in Basal stage, 1162 genes after PMSG stimulation and 1526 genes at 4h post-hCG. A list of the DEGs in *Erβ*^null^ GCs compared to wildtype at Basal, PMSG, and hCG 4h time points are presented in **Supplementary Tables 7-9**.

### 3.3. ERβ-regulated genes in GCs 48h post-PMSG

Comparison of the PMSG group to the unstimulated (Basal) group within the same genotype exhibited differential expression of 1720 (591 + 1129) genes in wildtype GCs, and 1085 (447 + 638) genes in *Erβ*^null^ GCs. Direct comparison of PMSG-treated *Erβ*^null^ GCs to PMSG-treated wildtype GCs identified a total of 1162 DEGs with 453 genes showing downregulation and 709 showing upregulation. These results show a wide divergence in the gene expression profiles in GCs from wildtype and *Erβ*^null^ rat ovaries. In the **Fig. 3A** Venn diagram, we compared the PMSG-upregulated (from Basal) genes in wildtype GCs (total 591) and *Erβ*^null^ GCs (total 447) to those DEGs (total 453) that were downregulated in PMSG-treated *Erβ*^null^ GCs compared to wildtype GCs. In the side by side comparison of *Erβ*^null^ and wildtype GCs, only 201 (161+40) genes showed a similar pattern in response to PMSG in both, while 431 (296+135) genes were upregulated in wildtype GCs only and 246 (238+8) genes in *Erβ*^null^ GCs only. Remarkably, **175** (135+40) genes which were upregulated in wildtype and *Erβ*^null^ GCs after PMSG treatment were downregulated in *Erβ*^null^ GCs when compared to wildtype PMSG-treated GCs (**Fig. 3A**).

**Figure 3.**
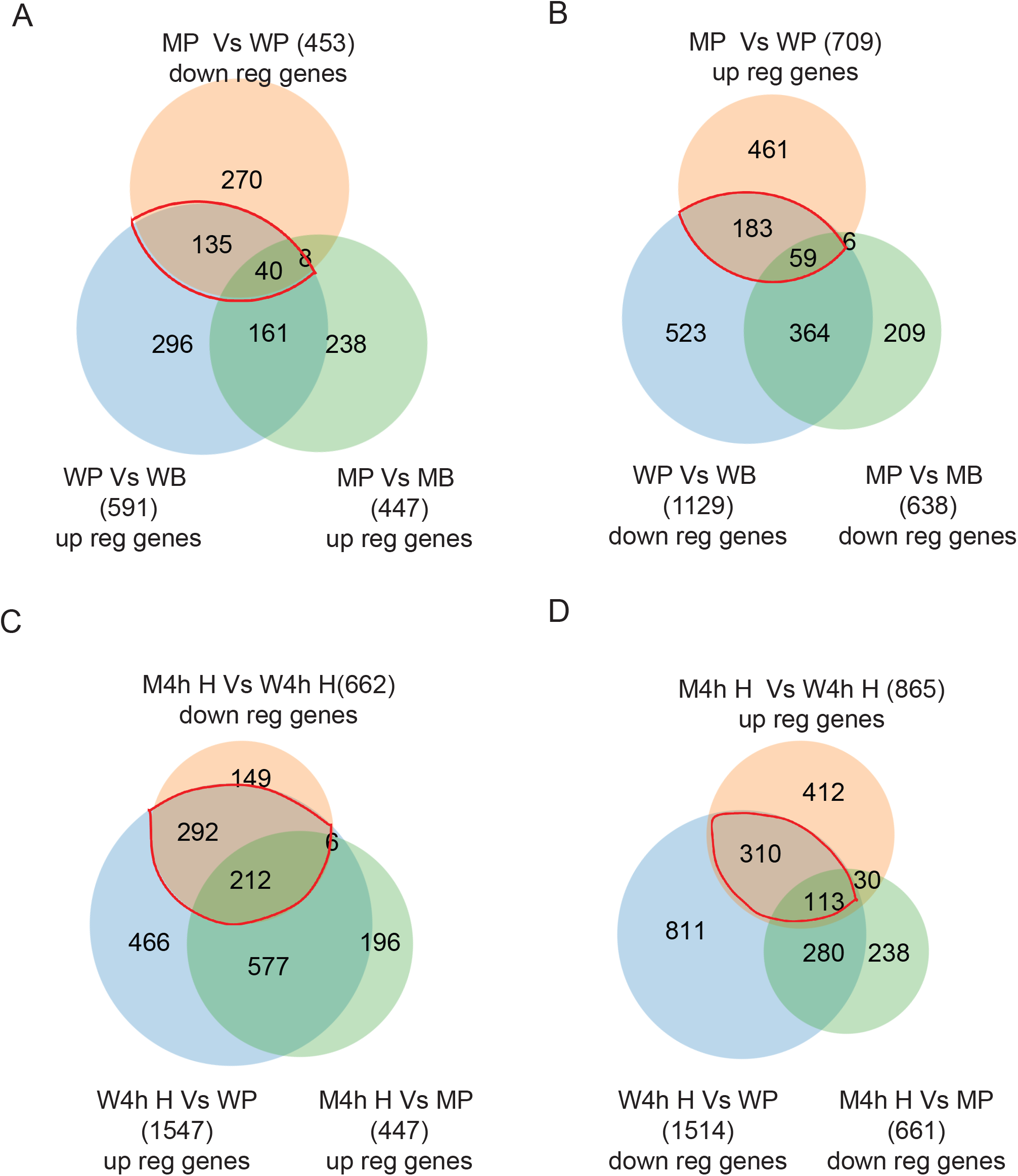
Identification of PMSG- and hCG-regulated genes that are dependent on ERβ signaling. Upregulated (up reg) genes from the Basal to PMSG treated group in wildtype (WP vs WB) granulosa cells (GCs) were compared with that of *Erβ*^*null*^ GCs (MP vs MB). These Venn diagrams were overlapped with the genes that were differentially downregulated (down reg) in PMSG-treated *Erβ*^*null*^ GCs compared to that of wildtype GCs (MP vs WP) to identify the potential PMSG-induced genes that are dependent on ERβ signaling (**A**). Downregulated genes from the Basal to PMSG group in wildtype GCs (WP vs WB) were compared with that of *Erβ*^*null*^ GCs (MP vs MB). These comparisons were overlapped with the genes that were upregulated in PMSG-treated *Erβ*^*null*^ GCs (MP vs WP) to identify the PMSG-inhibited genes that are dependent on ERβ (**B**). Similarly, upregulated genes in PMSG to hCG 4h wildtype GCs (W4h H vs WP) were compared to that of *Erβ*^*null*^ GCs (M4h H vs MP), which were overlapped with the genes downregulated in *Erβ*^*null*^ GCs 4h after hCG treatment (M4hH vs W4h H), to identify the hCG-induced genes that are dependent on ERβ (**C**). In addition, downregulated genes from PMSG to hCG 4h wildtype GCs (W4h H vs WP) were compared to that of *Erβ*^*null*^ GCs (M4h H vs MP), and with the differentially upregulated genes in *Erβ*^*null*^ hCG 4h GCs (M4h H vs W4h H), to detect the hCG inhibited genes that are dependent on ERβ (**D**). WB, Wildtype Basal; WP, Wildtype PMSG; W4h H, Wildtype 4h post-hCG; MB, *Erβ*^*null*^ Basal; MP, *Erβ*^*null*^ PMSG; M4h H, *Erβ*^*null*^ 4h post-hCG,

In the **Figure 3B** Venn diagram, we compare the genes downregulated (from Basal) after PMSG treatment, total 1129 in wildtype and 638 in *Erβ*^null^ GCs, to 709 DEGs that were upregulated in PMSG-treated *Erβ*^null^ GCs compared to wildtype GCs. Again, the side by side comparison of *Erβ*^null^ and wildtype GCs indicated that only a small proportion of 423 (59+364) genes showed a similar pattern in response to PMSG in both, while 706 (523+183) genes were upregulated in wildtype GCs only and 215 (209+6) genes in *Erβ*^null^ GCs only (**Fig. 3B**). Remarkably, **242** (183+59) genes which were downregulated in wildtype and *Erβ*^null^ GCs were upregulated in *Erβ*^null^ GCs when compared to wildtype PMSG-treated GCs (**Fig. 3B**).

### 3.4. ERβ-regulated genes in GCs 4h post-hCG

Comparative analyses were performed with the 4h post-hCG GCs (**Fig 3C** and **D**) and their respective PMSG controls as described above for the post-PMSG group. While hCG treatment altered the expression of 3061 genes (1547 upregulated and 1514 downregulated) in wildtype GCs, expression of only 1108 genes (447 upregulated and 661 downregulated) were significantly modulated in *Erβ*^null^ GCs. Direct comparison of hCG-treated *Erβ*^null^ GCs to hCG-treated wildtype GCs indicated a total of 1527 DEGs, with 662 genes showing downregulation and 865 showing upregulation in the *Erβ*^null^ GCs. Of the total 1547 upregulated genes in wildtype hCG 4h GCs, 789 (577 +212) genes showed similar expression patterns in *Erβ*^null^ hCG 4h GCs (**Fig. 3C**). 758 (466 +292) upregulated genes were unique to wildtype GCs and 212 (196 + 6) to *Erβ*^null^ GCs. Remarkably, **504** (292+212) genes that were upregulated in wildtype hCG 4h GCs were downregulated in *Erβ*^null^ hCG 4h GCs (**Fig. 3C**).

Of the total 1514 genes downregulated in wildtype hCG 4h GCs, 393 (280+113) genes were also downregulated in *Erβ*^null^ hCG 4h GCs (**Fig. 3D**). 1121 (811+310) downregulated genes were unique to wildtype GCs and 268 (238 +30) to *Erβ*^null^ GCs (**Fig. 3D**). Furthermore, we identified **423** (310 +113) genes that were downregulated in wildtype hCG 4h GCs but were upregulated in *ERβ*^*null*^ hCG 4h GCs (**Fig. 3D**).

### 3.5. ERβ-regulated FSH/FSHR target genes

As PMSG predominantly acts on FSH receptors (FSHRs), we examined the PMSG-regulated genes using Ingenuity Pathway Analysis (IPA) specifically for FSH/FSHR signaling. IPA demonstrated that a large number of the **175** genes (**Fig. 3A**) that were upregulated by PMSG treatment (from Basal) in wildtype or *Erβ*^null^ GCs but downregulated in PMSG-treated *Erβ*^null^ GCs (compared to PMSG-treated wildtype) have roles in folliculogenesis **(Fig. 4A)**. Further analyses determined that these genes downregulated in *Erβ*^*null*^ GCs are specifically involved in steroid metabolism, follicle development, molecular transport, cell survival, ovulation, and fertility (**Fig. 4A**).

**Figure 4.**
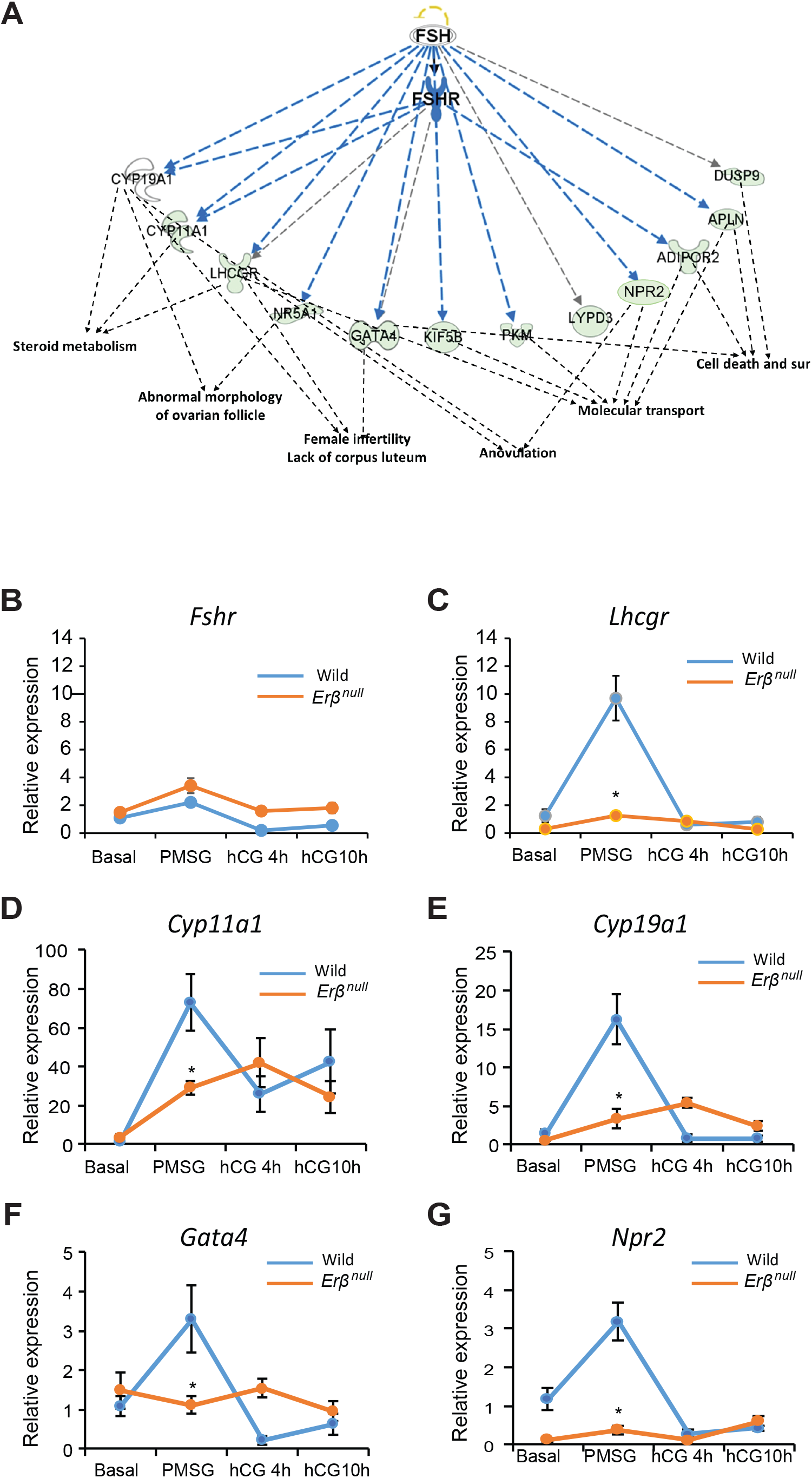
PMSG-regulated genes that are ERβ dependent and involved in folliculogenesis. PMSG target genes that were identified in the **Fig. 3A** were subjected to Ingenuity Pathway Analysis (IPA). IPA showed the group of downregulated genes that have FSH/FSHR as an upstream regulator (**A**) are involved in steroid metabolism, folliculogenesis, ovulation, molecular transport, cell death, and cell survival (sur). The expression of these genes was further validated by RT-qPCR analysis (**B-G)**. We did not detect any significant difference in the expression of *Fshr* between wildtype and *Erβ*^*null*^ GCs (**B**). However, we observed a marked downregulation of *Lhcgr* in *Erβ*^*null*^ GCs (**C**). In addition, *Erβ*^*null*^ GCs showed downregulation of steroidogenic enzymes *Cyp11a1* and *Cyp19a1* (**D, E**) as well as *Gata4* and *Npr2* (**F, G**). RT-qPCR data are represented as mean ± SEM. n ≥ 6. *P ≤ 0.05.

We observed that the expression of *Fshr* was not decreased in PMSG-treated *Erβ*^*null*^ GCs (**Fig. 4B**), but several other genes known to be upregulated by FSH/FSHR signaling failed to respond appropriately (**Fig. 4C-G**). Importantly, PMSG-induced upregulation of *Lhcgr* was absent in *Erβ*^*null*^ GCs (**Fig. 4C**). PMSG-induced upregulation of steroidogenesis-related genes including *Cyp11a1* and *Cyp19a1* was also absent in *Erβ*^*null*^ GCs (**Fig. 4D** and **E**). Expression of the transcriptional regulator, *Gata4*, and *Npr2*, failed to be induced by PMSG in *Erβ*^*null*^ GCs (**Fig. 4F** and **G**). Interestingly, expression levels of these genes were quite low in both wildtype and *Erβ*^*null*^ GCs at the Basal stage before gonadotropin administration, reaching peak levels 48h after PMSG injection but decreasing to Basal levels 4h after hCG injection (**Fig. 4C-G**).

### 3.6. ERβ-regulated LH/LHCGR target genes

hCG is known to act on the LH receptors (LHCGRs) expressed on GCs, and we have examined the hCG-regulated genes in IPA specifically for LH/LHCGR signaling. IPA identified that a large number of the **504** genes (**Fig. 3C**) that were significantly upregulated (from PMSG) in hCG-stimulated wildtype or *Erβ*^*null*^ GCs but downregulated in hCG-stimulated *Erβ*^*null*^ GCs (compared to hCG-stimulated wildtype). These genes are involved in preovulatory cumulus cell expansion, oocyte maturation, and the induction of ovulation (**Fig. 5A**).

**Figure 5.**
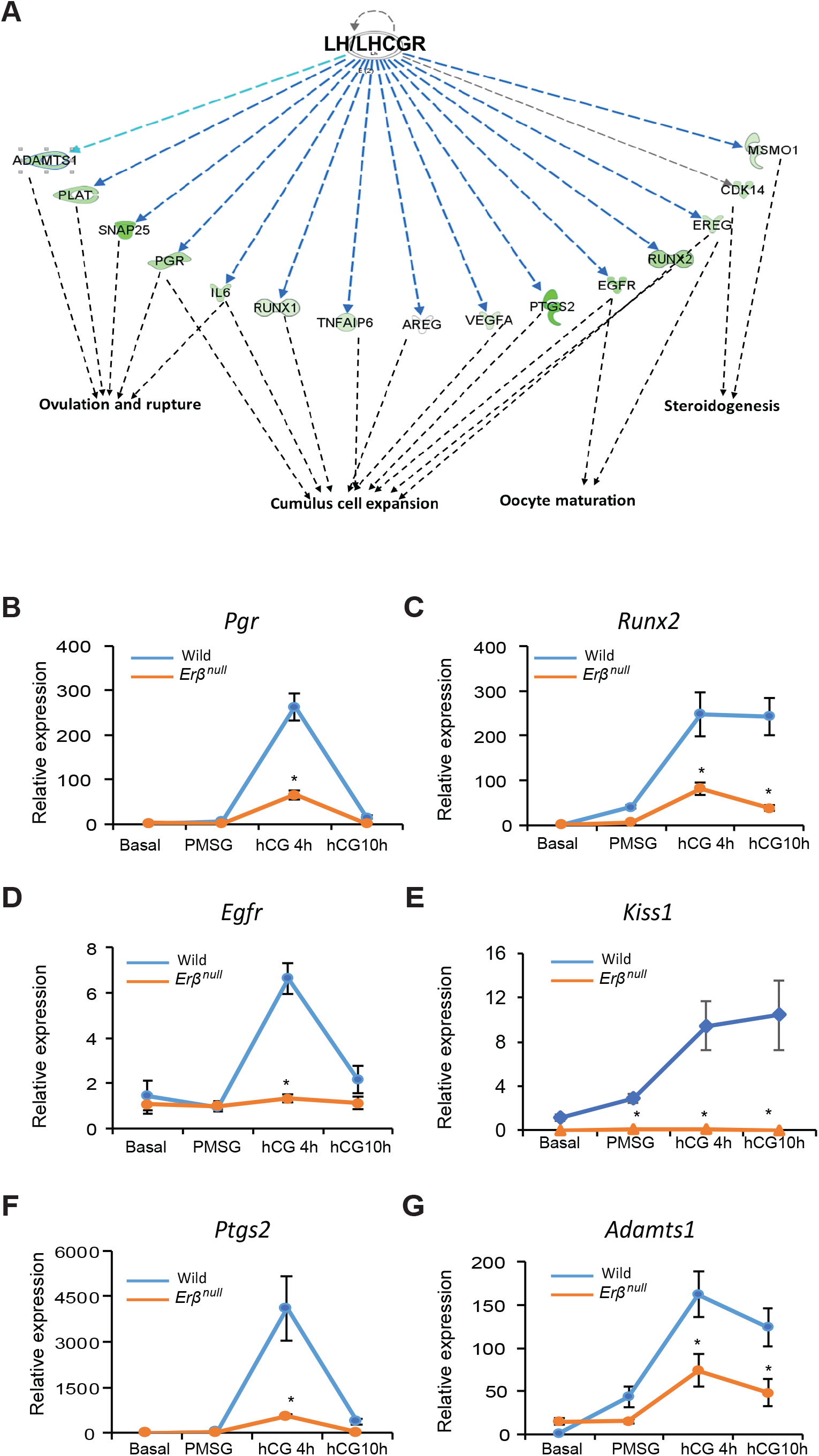
hCG-regulated genes that are dependent on ERβ and involved in follicle maturation. hCG-regulated genes that were identified in **Fig. 3C** were subjected to Ingenuity Pathway Analysis (IPA). IPA showed a subset of LH/LHCGR-regulated genes, which are involved in steroidogenesis, cumulus cell expansion, oocyte maturation, and ovulation (**A**). RT-qPCR analysis confirmed the expression data (**B-G**). hCG-induced genes involved in follicle maturation (i.e., *Pgr, Runx2, Egfr*, and *Kiss1*) (**B-E**) and induction of ovulation (i.e., *Ptgs2* and *Adamts1*) (**F, G**) were observed in wildtype GCs, which were not upregulated in *Erβ*^*null*^ GCs. RT-qPCR data are represented as mean ± SEM. n ≥ 6. *P ≤ 0.05.

In wildtype GCs, expression levels of transcriptional regulators *Pgr* and *Runx2* were low after PMSG treatment but increased rapidly after hCG treatment as expected (**Fig. 5B** and **C**). Administration of hCG failed to upregulate the expression of *Pgr* and *Runx2* in *Erβ*^*null*^ GCs, but not to the extent that was observed in wildtype GCs (**Fig. 5B** and **C**). Among the other key genes involved in preovulatory follicle development and ovulation, *Egfr* (**Fig. 5D**) and *Ptgs2* (**Fig. 5F**) showed expression patterns in *Erβ*^*null*^ GCs similar to that of *Pgr* in wildtype GCs (**Fig. 5B**). In contrast, *Kiss1* (**Fig. 5E**) and *Adamts1* (**Fig. 5G**) showed elevated expression in *Erβ*^*null*^ GCs similar to that of *Runx2* in wildtype GCs (**Fig. 5C**). hCG-induced upregulation of these genes (*Egfr, Ptgs2, Kiss1*, and *Adamts1*) was either blunted (**Fig. 5B, C, F** and **G**) or absent (**Fig. 5D** and **E**) in *Erβ*^*null*^ GCs.

### 3.7. ERβ-regulated novel genes involved in follicle development

We also identified novel gonadotropin-stimulated genes that were differentially upregulated in wildtype GCs but failed to respond in *Erβ*^*null*^ GCs. We speculate that these genes may contribute to gonadotropin-stimulated ovarian follicle development. However, IPA analyses did not link these genes to either gonadotropin or ERβ signaling. We have selected and validated the expression of theses novel genes based on their higher TPM values and relative fold changes.

Among these genes, *Jaml* (**Fig. 6A**), *Galnt6* (**Fig. 6B**), *Znf750* (**Fig. 6C**), and *Dusp9* (**Fig. 6D**) were upregulated by PMSG treatment and downregulated after hCG administration in wildtype GCs. Remarkably, the expression of *Jaml* (**Fig. 6A**), which is a cell adhesion molecule, was significantly higher at Basal stage in wildtype GCs. Two other novel genes, *Wnt16* (**Fig. 6E**) and *Mageb16* (**Fig. 6F**), were upregulated in wildtype GCs only after hCG treatment. While the expression of *Wnt16* (**Fig. 6E**) came down to Basal level 10h after hCG treatment, *Mageb16* (**Fig. 6F**) level remained elevated. These genes exhibited minimal changes in *Erβ*^*null*^ GCs after gonadotropin treatment.

**Figure 6.**
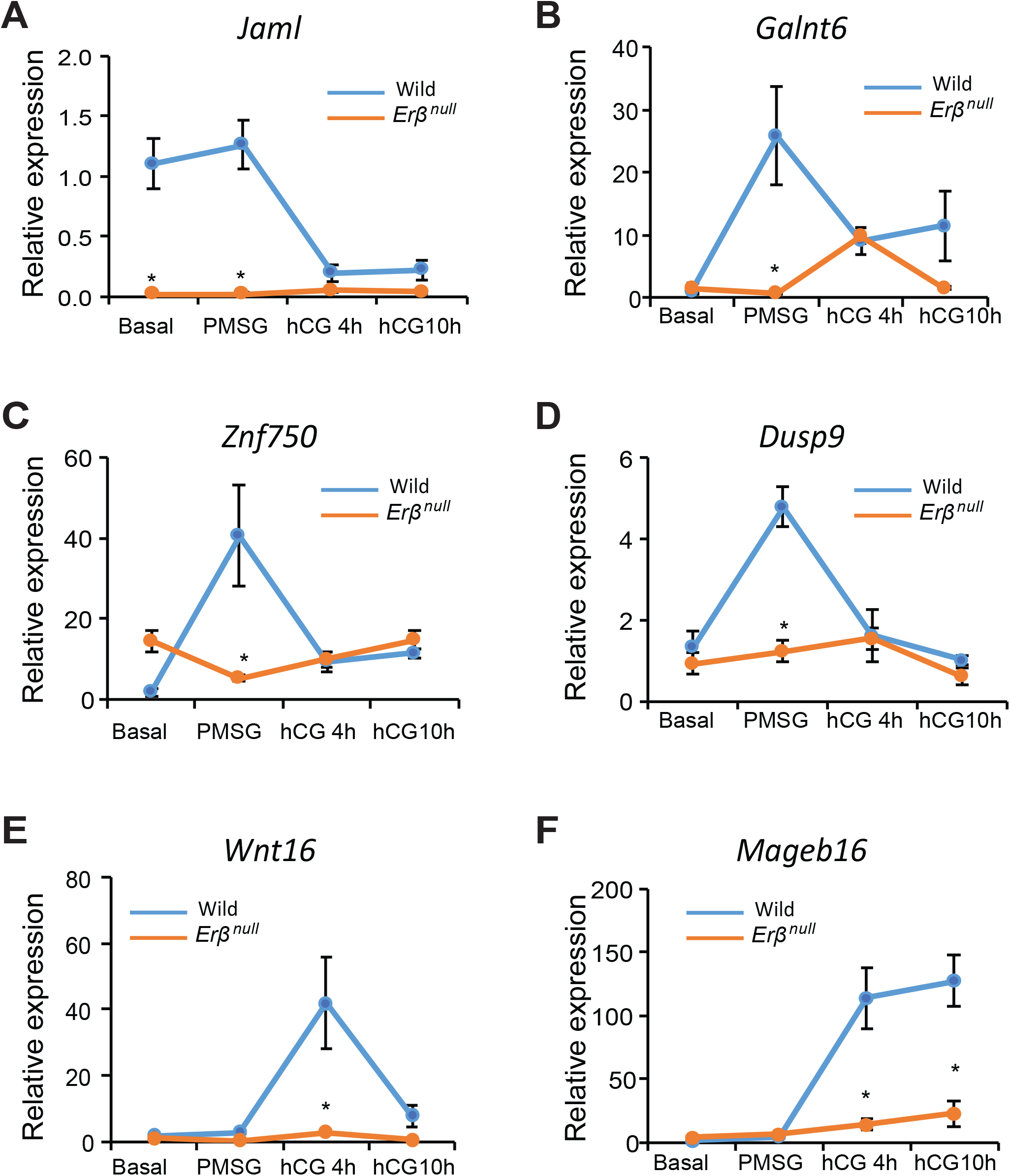
Gonadotropin-induced and ERβ-regulated novel genes with potential roles in follicle development. We identified several PMSG (**A-D**) and hCG (**E, F**) regulated genes, which were not previously shown as ERβ-regulated genes or regulated by gonadotropins. We identified these genes based on their relative abundance (TPM values), fold changes in *Erβ*^*null*^ GCs. PMSG induced genes included *Jaml, Galnt6, Znf750*, and *Dusp9* (**A-D**). hCG induced genes included *Wnt16* (**E**), and *Mageb16* (**F**). The expression of these genes was validated by RT-qPCR analysis (**A-F**). RT-qPCR data are represented as mean ± SEM. n ≥ 6. *P ≤ 0.05.

**Figure 7.**
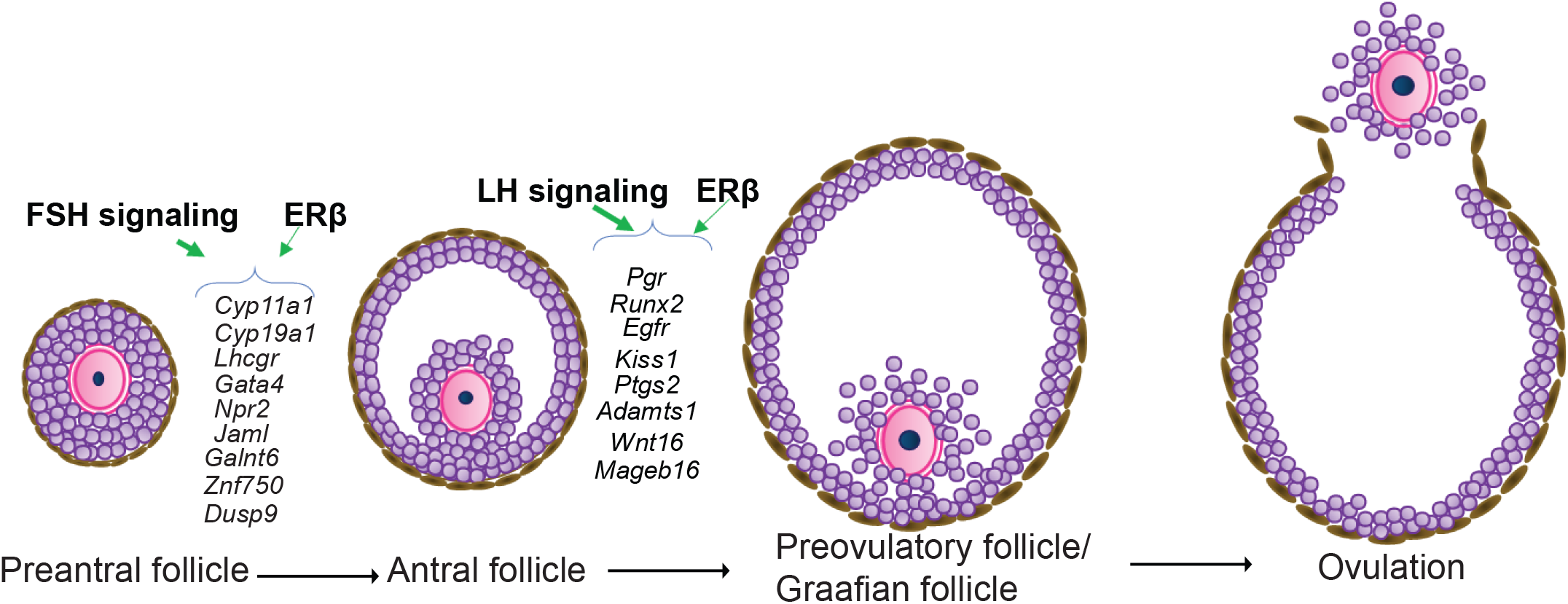
ERβ-regulated granulosa cell genes that are crucial for gonadotropin induced follicle maturation. Expression of FSH/FSHR-induced genes including *Cyp11a1, Cyp19a1, Lhcgr, Gata4, Npr2, Jaml, Galnt6, Znf750*, and *Dusp9* as well as expression of LH/LHCGR-induced genes including *Pgr, Runx2, Egfr, Kiss1, Ptgs2, Adamts1, Wnt16*, and *Mageb16* are dependent on ERβ-signaling. These genes play a crucial role in the preovulatory maturation of ovarian follicles.

## 4. DISCUSSION

*Erβ*^*null*^ female rodents are infertile due to defective follicle development and failure of ovulation (Couse and Korach, 1999; Couse et al., 2003; Rumi et al., 2017). In order to elucidate the specific defects in follicle development, we studied exogenous-gonadotropin-induced synchronized follicle development using 4-wk-old female rats. We observed that gonadotropin stimulation fails to initiate the selection and dominance of antral follicles in *Erβ*^*null*^ rats. The presence of many antral follicles of similar size and lack of preovulatory Graafian follicles suggest that ERβ is essential for follicle development beyond the antral stage.

ERβ is the major estrogen receptor in the ovary (Byers et al., 1997a; Couse et al., 2004b; Pelletier and El-Alfy, 2000b; Słomczyńska et al., 2001), and GCs are the major cell-type in ovarian follicles that express ERβ (Cheng et al., 2002; Paterni et al., 2014). ERβ plays an important role in the transcriptional regulation of GC genes that are involved in the GC-oocyte interactions required for the development of ovarian follicles (Couse et al., 2005b; Khristi et al., 2018; Rumi et al., 2017). In this study, we analyzed the GC transcriptome before and after gonadotropin administration to identify the ERβ-regulated genes (direct or indirect) that are involved in follicle maturation beyond the antral stage. Using a systematic approach, we identified the DEGs that are responsible for PMSG- and hCG-induced development of wildtype ovarian follicles. Next, among these genes, we identified DEGs that showed a similar or opposite expression pattern in *Erβ*^*null*^ GCs. This unique approach of differential transcriptome analysis enabled us to identify the ERβ-regulated genes that are dysregulated after PMSG or hCG administration in *Erβ*^*null*^ rats. These ERβ-regulated genes were further subjected to IPA to identify key genes involved in steroidogenesis, follicle development, cumulus expansion, oocyte maturation, and ovulation.

It is noteworthy that, for the transcriptome and subsequent IPA analyses, we have considered PMSG treatment to be representative of FSH/FSHR activation and hCG treatment of LH/LHCGR activation. Despite a normal level of *Fshr* expression in *Erβ*^*null*^ GCs, PMSG treatment failed to induce many of the FSH/FSHR target genes including *Lhcgr, Cyp11a1, Cyp19a1, Gata4*, and *Npr2*; however, expression of some other known targets like *Star* remained unaffected. This suggests that upregulation of a subset of PMSG-induced genes also requires the presence of ERβ in GCs. In wildtype GCs, a low level of expression before gonadotropin treatment (Basal stage) and marked upregulation after PMSG administration suggests that a high level of these proteins is necessary during this stage of follicle development or to execute the responses of LH/LHCGR signaling during the subsequent stage. Moreover, a sharp decrease of selective transcript levels after hCG administration suggests that either LH/LHCGR signaling leads to abrupt downregulation of transcription or degradation of those mRNAs. Such spatiotemporal patterns of expression in response to gonadotropins are characteristic of many of the GC genes involved in follicle maturation (Bédard et al., 2003; Chakravarthi et al., 2015; Chakravarthi et al., 2016; Chen et al., 2009; Lakshminarayana et al., 2014; Ronen-Fuhrmann et al., 1998). However, those reports were based on studying one or a few genes, but our study demonstrates the spatiotemporal pattern of gene expression among the whole transcriptome in wildtype and *Erβ*^*null*^ GCs.

PMSG stimulation failed to upregulate the expression of *Lhcgr* in *Erβ*^*null*^ GCs, which is essential for follicle development beyond the antral stage (LaPOLT et al., 1990; Lu et al., 1993; Richards and Ascoli, 2018). *Lhcgr* knockout mice exhibited an ovarian phenotype with normal development of primary, secondary, and antral follicles but a lack of preovulatory Graafian follicles (Lei et al., 2001), which emphasizes the importance of ERβ-regulated *Lhcgr* expression. *Cyp19a1* knockout mice also developed antral follicles that failed to ovulate (Fisher et al., 1998). It has been reported that decreased levels of *Cyp11a1* or *Cyp19a1* disrupt the development of antral and Graafian follicles (Gershon and Dekel, 2020). The reduced level of *Gata4* expression in *Erβ*^*null*^ GCs may contribute to reduced GC proliferation and theca cell recruitment, which would impair the maturation of follicles to the preovulatory stage (Bennett et al., 2013; Efimenko et al., 2013; LaVoie, 2014). Additionally, NPR2 plays an important role in GCs by inducing cGMP levels, which inhibits phosphodiesterase 3A, prevents cAMP hydrolysis, and retrieves the oocytes from meiotic arrest (Arroyo et al., 2020; Potter et al., 2006; Zhang et al., 2010).

We identified that LH/LHCGR target genes, particularly the expression of genes involved in cumulus cell expansion, steroidogenesis, oocyte maturation, and ovulation, were affected in *Erβ*^*null*^ GCs. This effect was in part due to the failure of PMSG-induced *Lhcgr* upregulation in *Erβ*^*null*^ GCs. We detected that hCG-induced upregulation of *Pgr* in GCs is also dependent on ERβ. Progesterone signaling is essential for follicle maturation and induction of ovulation (Jo et al., 2002; Kubota et al., 2016; Robker et al., 2009). Progesterone signaling regulates the expression of *Egfr, Ptgs2*, and *Adamts1* (Robker et al., 2000), which was affected in *Erβ*^*null*^ GCs. Studies have shown that EGFR signaling can regulate the expression of *Runx2* and *Plat*, induce production of steroids, and promote oocyte maturation (Jamnongjit et al., 2005). EGFR signaling also plays an essential role in the transition of antral follicles to the preovulatory stage (Panigone et al., 2008; Wu et al., 2019). RUNX2 regulates the expression of *Ptgs2* and *Runx1*, impacting the ovulatory process (Lee-Thacker et al., 2018; Liu et al., 2009). We observed an absence of *Ptgs2* upregulation in *Erβ*^*null*^ GCs, which might be due to the failure of *Runx2* induction in response to hCG stimulation. PTGS2 is involved in prostaglandin synthesis, preovulatory follicle maturation, and ovulation (Laurincik et al., 1993; Siddappa et al., 2015). We previously reported that the expression of *Kiss1* in GCs is induced by gonadotropins, which is dependent on ERβ signaling (Chakravarthi et al., 2018). Intraovarian administration of kisspeptin increased the number of preovulatory follicles and decreased the number of antral follicles, whereas treatment with kisspeptin antagonist KP234 had the opposite effect, suggesting a requirement for kisspeptin during preovulatory follicle maturation (Fernandois et al., 2016). Recently, we have also shown that GC-derived KISS1 plays a role in follicle development and oocyte maturation (Chakravarthi et al, 2020). In addition to *Ptgs2*, hCG-induced upregulation of *Adamts1* was not detected in *Erβ*^*null*^ GCs. It has been suggested that LH/LHCGR-induced ERK activation is regulated by ADAMTS1, a metalloprotease, which is also involved in cumulus expansion and induction of ovulation (Arroyo et al., 2020; Madan et al., 2003; Shozu et al., 2005; Yung et al., 2010).

We also identified a subset of gonadotropin-induced genes in GCs, which were dependent on ERβ, but they were not known to be regulated by gonadotropin or ERβ signaling or to be involved in ovarian follicle development. Among these novel genes, JAML is a cell-cell adhesion protein and engaged in homophilic interactions (Moog-Lutz et al., 2003). Expression of *Jaml* was high in the Basal and PMSG groups and decreased after hCG stimulation in wildtype, but, in *Erβ*^*null*^, its expression was very low in all stages. The high level of expression in the Basal and PMSG stages suggests that JAML might have a role in preantral follicles, especially in GC-GC or GC-oocyte interactions. GALNT6 has been shown to be essential for the activation of EGFR by phosphorylation and by modification of O-glycosylation (Lin et al., 2017). It is well known that LH-induced EGFR signaling plays a critical role in preovulatory follicle development by activating MAPK pathways and inducing the steroidogenic enzymes (Jamnongjit et al., 2005; Panigone et al., 2008; Wu et al., 2019). ZNF750 is a cell cycle regulator that acts by inducing *Klf4* gene expression (Boxer et al., 2014; Liu et al., 2019; Sen et al., 2012). KLF4 increases the susceptibility of GCs to apoptosis by downregulating *Bcl2* expression and promotes LH induced luteal transition of GCs (Choi and Roh, 2018). Dual-specific phosphatases (DUSPs) dephosphorylate MAPKs and thereby inhibiting their activities (Chen et al., 2019). We speculate that PMSG-induced *Dusp9* suppresses ERK activation to allow proper development of preantral follicles in wildtype ovaries. In contrast, in *Erβ*^*null*^ ovaries, the lack of DUSP9 results in aberrant activation of ERK and leads to premature formation of many small antral follicles. In this study, we also observed a failure of PMSG- or hCG-stimulated induction of either *Wnt16* or *Mageb16*. WNT16 is a positive regulator of the canonical WNT pathway and it plays a potential role in the dominant follicle selection (Gupta et al., 2014). Mage proteins interact with nucleic acids or target proteins and regulate different cell processes (Rajagopalan et al., 2011). *Mageb16* was reported to be a testis-specific gene (Liu et al., 2014), and the expression of *Mageb16* was found to be low in the ovary (Liu et al., 2014). Interestingly, we observed a drastic increase in the expression of *Mageb16* in wildtype GCs after gonadotropin injection, which proves that gonadotropin stimulation is necessary for the expression of *Mageb16* in ovaries. Based on the expression and its general function, *Mageb16* may have a role in the later stages of follicle development and further studies are needed to confirm its role.

Ours results demonstrate the spatiotemporal pattern of gonadotropin-induced gene regulation in GCs that is essential for ovarian follicle maturation. These datasets may serve as the reference for understanding gonadotropin regulation of GC genes. Although a group of gonadotropin-regulated genes showed similar expression patterns in both wildtype and *Erβ*^*null*^ GCs, others were unique to each genotype. We were particularly interested in a subset of gonadotropin-regulated genes characterized by a remarkably different expression pattern in *Erβ*^*null*^ GCs. These included FSH signaling regulated genes like *Lhcgr, Cyp11a1, Cyp19a1, Gata4, Npr2, Jaml, Galnt6, Znf750*, and *Dusp6* and LH regulated genes *Pgr, Egfr, Adamts1, Ptgs2, Kiss1, Runx2, Wnt16*, and *Mageb16* (**Fig.7**). We also identified several novel genes including *Jaml, Galnt6, Znf750, Dusp6, Wnt16*, and *Mageb16*, which are gonadotropin-induced but dependent on ERβ, however, further studies are required to confirm their involvement in ovarian follicle maturation.

## Acknowledgments

Generation of these datasets was supported by funding from the KUMC SOM, COBRE (P30 GM122731), and K-INBRE.

