## Supplemental Table 2 for "Granulosa cell genes that regulate ovarian follicle development beyond the antral stage: the role of estrogen receptor β"

**Supplementary Table 2. List of primers used in the qPCR assays**

| **Symbol** | **Reference mRNA** | **Forward Primer** | **Reverse Primer** | **Amplicon** |
| --- | --- | --- | --- | --- |
| *Fshr* | NM_199237.1 | 471F-ACTTGCCAGCTGTTCACAAG | 579R-AAACTCAGTCCCATGAAGGA | 109 bp |
| *Lhcgr* | NM_012978.1 | 555F-ATGCTTTCCAAGGGATGAAT | 657R-GAGATTAGAGTCGTCCCATT | 103 bp |
| *Cyp19a1* | NM_017085.2 | 794F-CCCATGGCAGATTCTTGTGG | 1009R-TGATGCCATTCTCGTGCATG | 216 bp |
| *Cyp11a1* | NM_017286.3 | 498F-CCCCTGACTCCATCAAGAAC | 697R-GTGGAACCTCTACGCTTGGT | 200bp |
| *Gata4* | NM_144730.1 | 1029F-CTCCTACTCCAGCCCCTACC | 1315R-GCCGGTTGATACCATTCATCT | 287bp |
| *Dusp9* | NM_001037973.1 | 123F-TTGGCACTAGCTGTGGTGAG | 244R-AGGCATGACCGACTCAGACT | 122bp |
| *Pgr* | NM_022847.1 | 1004F-AAGTCCCTTTTGCTCCACCT | 1128R-GTAAACTGGGAACCCGTCCT | 125bp |
| *Egfr* | NM_031507.1 | 602F-TTAGCAACAACCCCATCCTC | 779RTTCTGGCAGTTCTCCTCTCC | 178bp |
| *Runx2* | NM_001278483.1 | 896F-CGACAGCCCCAACTTCCTGT | 1149R-CTTGGGGAGGATTTGTGAAG | 254bp |
| *Plat* | NM_013151.2 | 654F-GTGAAGCCCTGGTGCTATGT | 805R-GAGGCCTTGGATGTGGTAAA | 152bp |
| *Adamts1* | NM_024400.2 | 949F -TTCCACATCCTGAGGCGAAG | 1150R-GCTGGACACAAATCGCTTCT | 202 bp |
| *Ptgs2* | NM_017232.3 | 237F-CCACTTCAAGGGAGTCTGGA | 415R GAAGGGCCCTGGTGTAGTAG | 179 bp |
| *Jaml* | XM_236198.9 | 249F-ATGCTTTGCCTCCTGACACT | 447R-GCTCCCCTTTCGAGAATACC | 199bp |
| *Galnt6* | NM_001172063.2 | 422F-GCTTTAATGCCTTTGCCAGT | 570R-CCAGGCTTCGTTGTGGAATA | 149bp |
| *Znf750* | XM_006247958.3 | 1436F-GCCTTCAAACCTGTCCAGAG | 1664R-CATAAGTGGCTGCAGGGT TT | 229bp |
| *Npr2* | NM_053838.1 | 834F-CCTTGATGTCTTTGGGGAGA | 1113R-GACTTGGGCATAGAGCAGGA | 280bp |
| *Kiss1* | NM_181692.1 | 26F-TGCTGCTTCTCCTCTGTGTG | 174R-AGGCTTGCTCTCTGCATACC | 149bp |
| *Adcyap1* | NM_016989.2 | 419F-TTACGATCAGGACGGAAACC | 590R-CAGCTGGTCCAAGACTTTGC | 172bp |
| *Wnt16* | NM_001109223.1 | 832F-TACCCGTCAGAAACACCACA | 982R-CAGGTTTTCACAGCACAGGA | 151bp |
| *Mageb16* | NM_001025013.1 | 66F-TGAGCTGTTGCCTTTGATTG | 256R-GGAACATGGCTCCTCAACAT | 191bp |
| *Rn18s* | NR_046237.1 | 1622F-GCAATTATTCCCCATGAACG | 1744R GGCCTCACTAAACCATCCAA | 123 bp |
