## Supplemental Table 1 for "Granulosa cell genes that regulate ovarian follicle development beyond the antral stage: the role of estrogen receptor β"

| **Genotype** | **Group** | **Study ID** | **Sample ID** | **Experiment** | **Run** | **Run Name** |
| --- | --- | --- | --- | --- | --- | --- |
| Wildtype | Basal | PRJNA551764 | SAMN12165660 | SRX6376732 | SRR9613662 | WB1_1.fq.gz |
| Wildtype | Basal | PRJNA551764 | SAMN12165661 | SRX6376731 | SRR9613663 | WB2_1.fq.gz |
| Wildtype | Basal | PRJNA551764 | SAMN12165662 | SRX6376734 | SRR9613660 | WB3_1.fq.gz |
| Wildtype | PMSG | PRJNA551764 | SAMN12165663 | SRX6376733 | SRR9613661 | WP1_1.fq.gz |
| Wildtype | PMSG | PRJNA551764 | SAMN12165664 | SRX6376728 | SRR9613666 | WP2_1.fq.gz |
| Wildtype | PMSG | PRJNA551764 | SAMN12165665 | SRX6376727 | SRR9613667 | WP3_1.fq.gz |
| Wildtype | hCG 4h | PRJNA551764 | SAMN12165666 | SRX6376730 | SRR9613664 | WG1.FCHTYFKBBXX_L4_IGTGGCC.bam |
| Wildtype | hCG 4h | PRJNA551764 | SAMN12165667 | SRX6376729 | SRR9613665 | WG2.FCHTYFKBBXX_L4_IGTTTCG.bam |
| Wildtype | hCG 4h | PRJNA551764 | SAMN12165668 | SRX6376735 | SRR9613659 | WG3.FCHTYFKBBXX_L4_ICGTACG.bam |
| ERβ^null^ | Basal | PRJNA551766 | SAMN12165688 | SRX6376754 | SRR9613684 | DB1_1.fq.gz |
| ERβ^null^ | Basal | PRJNA551766 | SAMN12165689 | SRX6376753 | SRR9613685 | DB2_1.fq.gz |
| ERβ^null^ | Basal | PRJNA551766 | SAMN12165690 | SRX6376756 | SRR9613682 | DB3_1.fq.gz |
| ERβ^null^ | PMSG | PRJNA551766 | SAMN12165691 | SRX6376755 | SRR9613683 | DP1_1.fq.gz |
| ERβ^null^ | PMSG | PRJNA551766 | SAMN12165692 | SRX6376750 | SRR9613688 | DP2_1.fq.gz |
| ERβ^null^ | PMSG | PRJNA551766 | SAMN12165693 | SRX6376749 | SRR9613689 | DP3_1.fq.gz |
| ERβ^null^ | hCG 4h | PRJNA551766 | SAMN12165694 | SRX6376752 | SRR9613686 | DG1.FCHTYFKBBXX_L4_IGAGTGG.bam |
| ERβ^null^ | hCG 4h | PRJNA551766 | SAMN12165695 | SRX6376751 | SRR9613687 | DG2.FCHTYFKBBXX_L4_IACTGAT.bam |
| ERβ^null^ | hCG 4h | PRJNA551766 | SAMN12165696 | SRX6376757 | SRR9613681 | DG3.FCHTYFKBBXX_L4_IATTCCT.bam |

**Supplementary Table 1. RNA-sequencing data-sets available in SRA website**

PRJNA551764 datasets are available at <https://www.ncbi.nlm.nih.gov/sra/?term=PRJNA551764>

PRJNA551766 datasets are available at <https://www.ncbi.nlm.nih.gov/sra/?term=PRJNA551766>
